# High-Resolution Cryo-EM Reveals Dynamics in the Murine Norovirus Capsid

**DOI:** 10.1101/693143

**Authors:** Joseph S. Snowden, Daniel L. Hurdiss, Oluwapelumi O. Adeyemi, Neil A. Ranson, Morgan R. Herod, Nicola J. Stonehouse

## Abstract

Rather than acting as rigid symmetrical shells to protect and transmit their genomes, the capsids of non-enveloped, icosahedral viruses co-ordinate multiple, essential processes during the viral life-cycle, and undergo extensive conformational rearrangements to deliver these functions. Capturing conformational flexibility has been challenging, yet could be key in understanding and combating infections that viruses cause. Noroviruses are non-enveloped, icosahedral viruses of global importance to human health. They are a common cause of acute non-bacterial gastroenteritis, yet no vaccines or antiviral agents specific to norovirus are available. Here, we use cryo-electron microscopy to study the high-resolution solution structures of infectious, inactivated and mutant virions of murine norovirus (MNV) as a model for human noroviruses. Together with genetic studies, we show that the viral capsid is highly dynamic. While there is little change to the shell domain of the capsid, the protruding domains that radiate from this are flexible and adopt distinct states both independently and synchronously. In doing so the viral capsid is able to sample a defined range of conformational space, with implications for the maintenance of virion stability and infectivity. These data will aid in developing the first generation of effective control measures against this virus.

## Introduction

High-resolution structural information has been key in improving our understanding of viral lifecycles. However, viral capsids commonly undergo profound conformational changes during their infection cycles, as well as more subtle dynamics that can be challenging to capture. Understanding such conformational changes can hold the key for the design and development of antiviral agents and vaccines, which are needed for many viral diseases, including those caused by noroviruses.

Noroviruses are globally prevalent pathogens that cause approximately 200,000 deaths each year in low- and middle-income countries^1^. An effective vaccine or antiviral agent against norovirus would provide significant health and economic benefits, but none have been approved to date. Although several clinical trials of virus-like particle (VLP) vaccines have been undertaken, results have been disappointing^2,3^. This is partly because the VLPs do not provide a sufficiently long-lived and appropriate immune response. It is therefore possible that the VLPs are not acting as an appropriate surrogate for the intact virion.

As members of the *Caliciviridae* family, norovirus virions consist of a ~7.5 kb positive-sense single-stranded RNA genome linked at its 5’ end to a viral protein (VPg). This genomic RNA is enclosed within a non-enveloped, *T = 3* capsid, approximately 40 nm in diameter^4,5^. While there have been several high-resolution reconstructions of VLPs reported previously, structures of infectious noroviruses have been limited to a resolution of 8.0 Å^5–9^. The capsid is formed from 90 dimers of the major structural protein, VP1, in one of three quasi-equivalent conformational states; A-type VP1 proteins make up the icosahedral five-fold axes and form AB dimers with B-type VP1 proteins, whereas C-type VP1 proteins form CC dimers at two-fold axes. Each VP1 monomer comprises an N-terminal region, a shell (S) domain, and a protruding (P) domain (subdivided into proximal (P1) and distal (P2) subdomains). The S domains interact to completely enclose the contents of the capsid within an icosahedral shell. P domains extend outward from this shell and mediate interactions with receptor molecules and co-factors, such as bile acids^10–12^. Located within the capsid interior are a limited number of molecules of the minor structural protein, VP2^13^. While VP2 is unresolved in all norovirus structures to date, structural studies with the related *Vesivirus*, feline calicivirus, showed that VP2 can form a portal-like assembly as a result of conformational changes caused by VP1 receptor engagement^14^. In doing so it is believed to play an important role in viral genome release into the target cell.

Although the aforementioned structural studies have provided a wealth of information, there is still little knowledge on the dynamic nature of the viral capsid. Previous studies have observed subtle changes within the P domains as well as dramatically different P domain conformations in different noroviruses and related *Caliciviruses* ^6,7^. However, this is complicated by several factors, not least that different morphologies have only been observed between different norovirus species rather than within a single species^15^. Furthermore, there is no structural information on infectious human noroviruses due to difficulties in culturing the virus and the only structure of an infectious norovirus particle (that of murine norovirus) demonstrates gross morphological differences compared to most human norovirus VLPs, including major rearrangements of the P domain dimers^6,7^.

In this study, we present the high-resolution solution structure of an infectious norovirus, which shows dramatic structural differences to a previously published reconstruction at lower resolution^6^. Our reconstruction is remarkably similar to the overall structure of most human norovirus VLPs, suggesting that the previous murine norovirus structure may capture a non-native or alternative conformational state. We then used a murine norovirus (MNV) reverse genetics system to study the stability and conformational flexibility of infectious norovirus, using *in vitro* evolution to generate a mutant virus with increased stability. Our analysis reveals that P domains are independently mobile elements that can sample a wide conformational space whilst maintaining infectivity. We hypothesise that this allows noroviruses to interact with a range of receptor molecules or co-factors and could improve antibody evasion. These are powerful selective advantages for viral growth and challenge the idea of viral capsids as static containers for their genomic RNA. This may pave the way for new ideas to generate better immune responses for vaccination or antiviral strategies.

## Results

### The cryo-EM reconstruction of wtMNV reveals dynamic P domains

Currently, the best-resolved 3D reconstruction of wild-type MNV (wtMNV) has a nominal resolution of 8.0 Å^6^, which is not sufficient to resolve the secondary structure of the VP1 fold. Here, we used cryo-EM to determine the structure of wtMNV at 3.1 Å resolution (Figure 1a, S1a). wtMNV was cultivated in RAW264.7 cells, then purified by ultracentrifugation through a sucrose cushion and two sucrose gradients, before being applied to lacey carbon grids and vitrified. Cryo-EM data collection parameters are given in Table S1.

**Figure 1.**
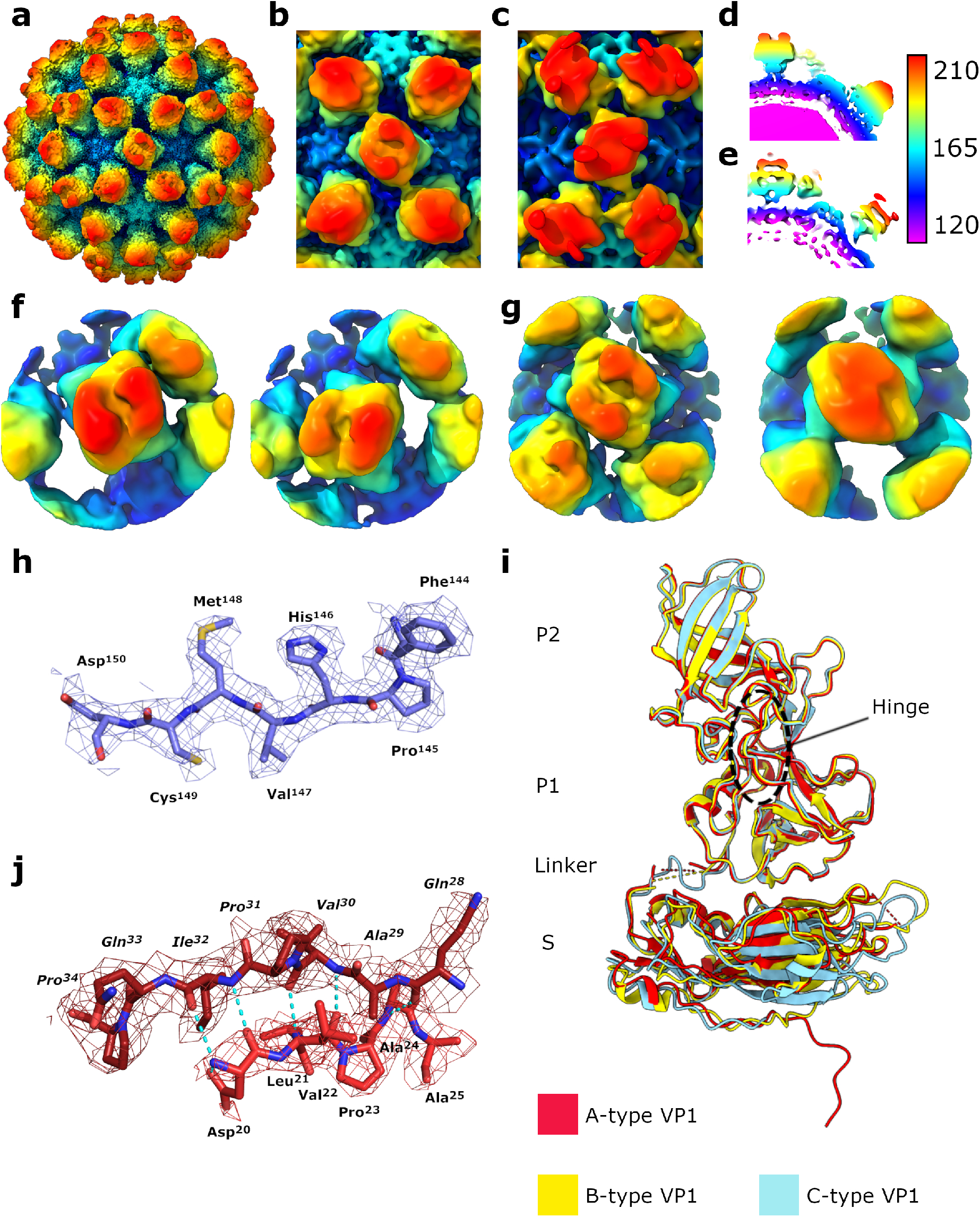
The 3.1 Å structure of wtMNV solved by cryo-EM. **(a)** Isosurface representation of the 3.1 Å wtMNV density map, shown at 1 σ and coloured according to the radial colouring scheme shown (Å). **(b,c)** Enlarged views centred on the icosahedral two-fold axis of **(b)** our wtMNV reconstruction, or **(c)** the previously published 8.0 Å cryo-EM reconstruction of wtMNV from Katpally *et al*. (2010)^21^, both low-pass filtered to 8.0 Å resolution, contoured to 1 σ and coloured according to the radial colouring scheme shown (Å). **(d,e)** Central sections through the reconstructions of wtMNV shown in (b) and (c), respectively. **(f,g)** Example class averages from focussed classification of **(f)** AB-type or **(g)** CC-type P domain dimers. **(h)** Representative EM density containing the refined atomic coordinates of VP1. **(i)** Overlaid atomic coordinates for different quasi-equivalent copies of MNV VP1, shown by the colouring scheme. The black dashed ring shows the hinge region connecting P2 to the C-terminal section of P1. **(j)** Atomic coordinates for N-terminal regions of adjacent A-type VP1 molecules, with polar contacts shown with blue dashed lines.

As expected, the MNV capsid shows *T* = 3 icosahedral symmetry, and is formed of ninety dimers of VP1. Interestingly, there are striking differences in the position and orientation of VP1 P domains relative to S domains, compared to the previously published 8.0 Å resolution cryo-EM reconstruction^6^ (Movie S1). The P domains from AB-type dimers (surrounding the icosahedral five-fold axes) are rotated approximately 90° anti-clockwise relative to their position in the previous structure, while the P domains from CC-type dimers (at icosahedral two-fold axes) are rotated approximately 70° anti-clockwise (Figure 1b,c). The P domains are also located much closer to the shell of S domains (separated by ~6 Å rather than ~16 Å), (Figure 1d,e).

While the S domains of VP1 are well resolved, density corresponding to the P domains had lower resolution (Figure S2a), likely reflecting their increased mobility. Reconstruction of a 3D density map from cryo-EM data involves the averaging of many images – if P domains are not rigidly held in the same position relative to the S domains, this averaging results in a blurring of density and loss of high-resolution information. The P2 subdomain, which is furthest away from the highly rigid S domain, is particularly poorly resolved, limiting the regions of the P domain that can be modelled. This is especially problematic for AB dimers, suggesting that they are more mobile than CC dimers.

In an attempt to address the lower resolution of the P domains, we performed 3D classification on the wtMNV dataset. Global 3D classification separated particles with better-resolved P domains (which were taken forward for the final icosahedral reconstruction) from those with extremely poorly-resolved P domains (which were excluded from the final reconstruction). However, this approach did not reveal a class with a distinct, alternative capsid conformation that might explain the diffuse nature of P domain density. This suggests that the lower resolution in this part of the map is a result of individual P domain dimer mobility, and not the coordinated movement of P domains across entire capsids.

With this observation in mind, we performed focussed 3D classification on AB- and CC-type P domain dimers separately (Figure S3a,b). This approach involves the assignment of sixty symmetrically redundant orientations to each particle and application of a mask to focus classification on a substructure within the virion – here, an AB- or CC-type P domain dimer. This approach revealed a striking diversity in the orientation and location of AB dimers (Figure 1f), but much less variation in CC dimer positioning (Figure 1g). This rationalises the lower resolution of A- and B-type P domain density than C-type P domain density described above. Complete capsids reconstructed from individual classes showed well-resolved density for the P domain dimer that was contained within the mask during focussed classification, but lower quality density for P domain dimers outside of this mask (Figure S3c). Importantly, these data are consistent with P domain dimers being mobile elements that move independently of each other on the capsid surface. However, it should be noted that this approach did not lead to a significant improvement in the quality of P domain density, presumably because the number of particles contributing to the final reconstruction was split between multiple classes, reducing the data in any individual reconstruction.

### An atomic model of wtMNV VP1

The quality of the data allowed us to build a hybrid atomic model for VP1 (Figure S4, Table S2). To construct an initial atomic model for refinement into our EM density map, a homology model of an MNV VP1 S domain was generated using the Phyre2 server^16^, based on the crystal structure of the Norwalk virus capsid (PDB: 1IHM)^9^. This homology model was rigid-body fitted into our density map and copies were fitted into each quasi-equivalent position (A, B and C) within the asymmetric unit, before each was manually edited and then refined to improve fit against the experimental cryo-EM density and the geometry of the model. The resolution in the S domain was sufficient to allow confident building of most residues up to the flexible linker region (Figure 1h). As expected, the flexible linker connecting the S and P domains was not resolved for A- or B-type VP1. However, the flexible linker for C-type VP1 was resolved, suggesting it is less mobile than the A- or B-type conformer, and correlating with the improved resolution of C-type P domains. To model the P domains, a crystal structure of an MNV VP1 P domain (PDB: 6C6Q)^11^ was fitted into each quasi-equivalent position, then refined against the cryo-EM density in combination with the S domain model, with secondary structure restraints imposed.

There are subtle but important differences between VP1 molecules occupying A-, B-and C-type quasi-equivalent positions (Figure 1i). RMSD values between the subset of residue pairs used for an alignment of the atomic coordinates were small (0.87 Å [A-B], 0.79 Å [A-C], 0.75 Å [B-C]), but increased when all residue pairs were included (2.77 Å [A-B], 2.46 Å [A-C], 2.35 Å [B-C]). This reflects a high degree of overall similarity between VP1 molecules, but with certain regions showing substantial variability. In particular, there are striking differences between the N-terminal regions of the three quasi-conformers. Most of the C-type VP1 N-terminal region is disordered (residues 1 – 29), whereas A- and B-type VP1 N-terminal regions are better resolved (missing residues 1 – 19 and 1 – 16, respectively) and have distinct conformations. The B-type N-terminal region remains close to the underside of the S domain and turns to run towards the icosahedral three-fold axis. Comparatively, the A-type N-terminal regions protrude deeper into the capsid to interact with adjacent A-type N-terminal regions around the icosahedral five-fold axis (Figure 1j).

In summary, these data show both the defined nature of VP1 quasi-conformers and the dynamic nature of the P domains.

### Thermal inactivation of wtMNV results in non-infectious but intact virus particles

Given the dynamic nature of the virion, we next aimed to capture *in vitro* defined conformations of the MNV virion that reflect the conformational changes a norovirus capsid undergoes during the viral life-cycle. Thermally stressing enteroviruses is an established approach to inducing alternative capsid conformations that are informative of those that occur during cell entry^17,18^. We therefore set out to identify structural changes that occur to the norovirus capsid after thermal stress. Our simple hypothesis was that more mobile elements of the viral capsid would be the first to change conformation upon heating.

This first required us to characterise the thermal stability of MNV virions. We therefore performed TCID_50_ assays with MNV after heating on a 30-second constant temperature ramp to identify a point at which the virus had lost >99.9% titre (Figure 2a). We also investigated capsid stability by performing PaSTRy assays, which employ two fluorescent dyes, SYTO-9 (which binds to nucleic acids) and SYPRO-Orange (which binds to hydrophobic regions of proteins), to assess the stability of viral capsids, independent of viral infectivity^19^ (Figure 2b). While there was a 10,000-fold reduction in infectivity at 61°C, PaSTRy assay data suggested that capsids remained essentially intact up to ~64°C, as minimal SYTO-9 fluorescence suggests that the viral RNA was not exposed to bulk solvent below this temperature. Confirming this, MNV heated to 61°C (which we termed heat inactivated MNV, or hiMNV) was incubated with RNase and no digestion of the RNA genome was observed (Figure 2c). Thus, we had identified a temperature at which the capsids had become irrevocably non-infectious, but were not disassociated into their component parts.

**Figure 2.**
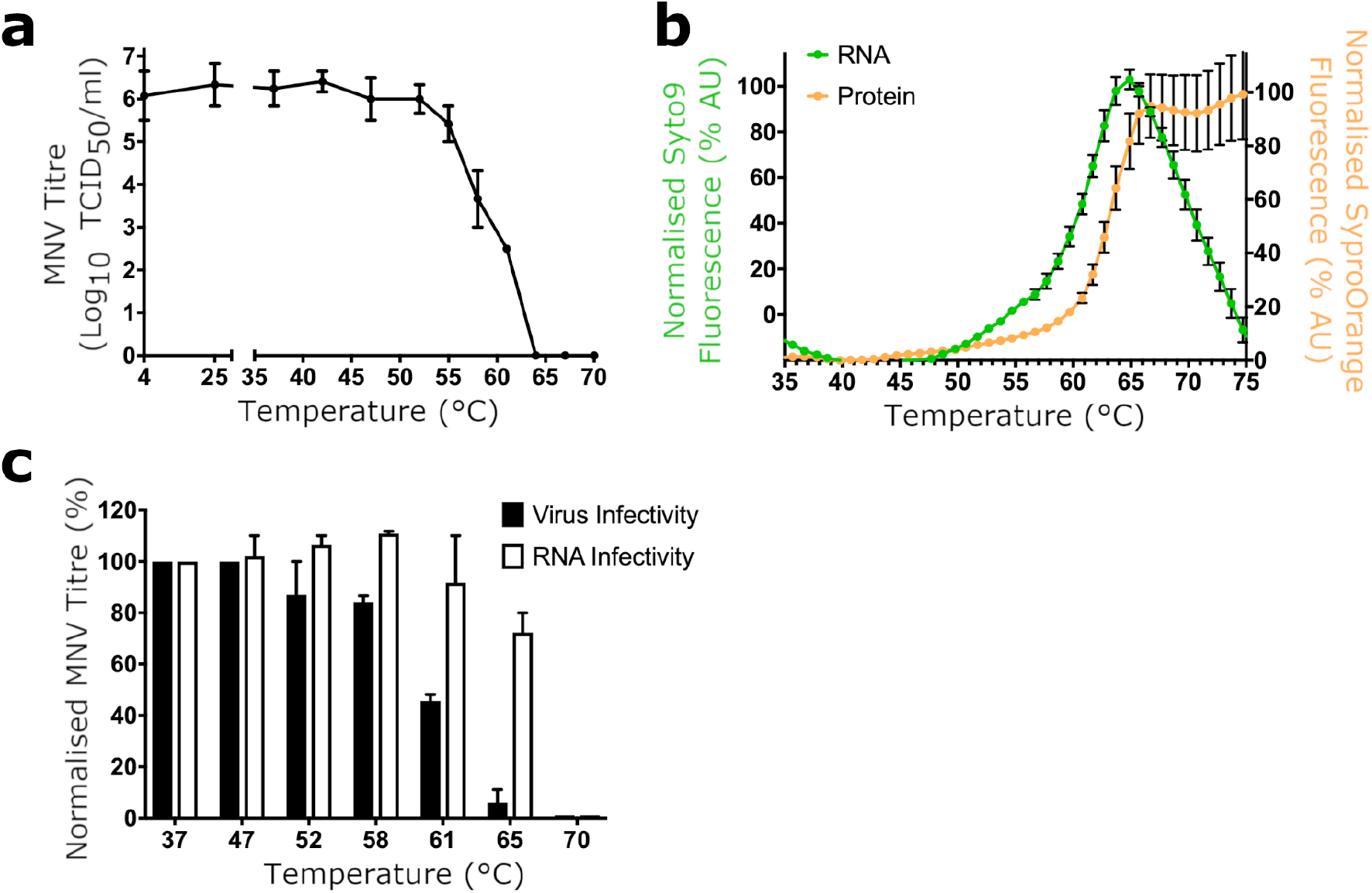
Thermal inactivation of wtMNV. **(a)** Samples of wtMNV were incubated at a range of temperatures up to 70°C on a 30 second constant temperature-ramp, before being immediately cooled on ice. Titres were determined by TCID_50_ assay on RAW264.7 cells (*n*=2 ± SEM). **(b)** wtMNV was purified by sucrose density gradient, dialysed into PBS and used for PaSTRy thermal stability assays using the nucleic acid dye SYTO-9 (green) and the protein dye SYPRO-Orange (orange) on a 30 second constant temperature-ramp (*n*=2 ± SD). **(c)** Samples of MNV were heated to the indicated temperatures, treated with RNase A and then titrated by TCID_50_ assay on RAW264.7 cells (n=2 ± SEM) or used to extract total RNA. The extracted RNA was transfected into BHK cells (which only permit a single round of replication), and resultant virus was harvested and titrated by TCID_50_ assay on RAW264.7 cells (*n*=2 ± SEM).

### The cryo-EM reconstruction of hiMNV reveals an increase in P domain mobility

To understand the structural changes that occurred during thermal stress, we determined the structure of hiMNV by cryo-EM at 2.9 Å resolution. This resolution is significantly higher than for wtMNV (2.9 Å vs 3.1 Å; Figure S1b, S2b), although considerably more data was available for the hiMNV reconstruction. There were no gross morphological differences in the positioning of the P domains with respect to the S domains compared to the wtMNV map.

Furthermore, while the strength of the S domain density was similar between the two maps, P domain density was weaker for hiMNV than for wtMNV (Figure 3a). This was also the case after applying a low-pass filter to limit the resolution of both the wtMNV and hiMNV reconstructions to 8.0 Å resolution, allowing a direct comparison with the previously published cryo-EM reconstruction of wtMNV^6^ (Figure 3b). This effect could result from either the fold of the P domain itself becoming less rigid, or the P domains becoming more mobile, and thus occupying a larger range of positions relative to the S domain shell.

**Figure 3.**
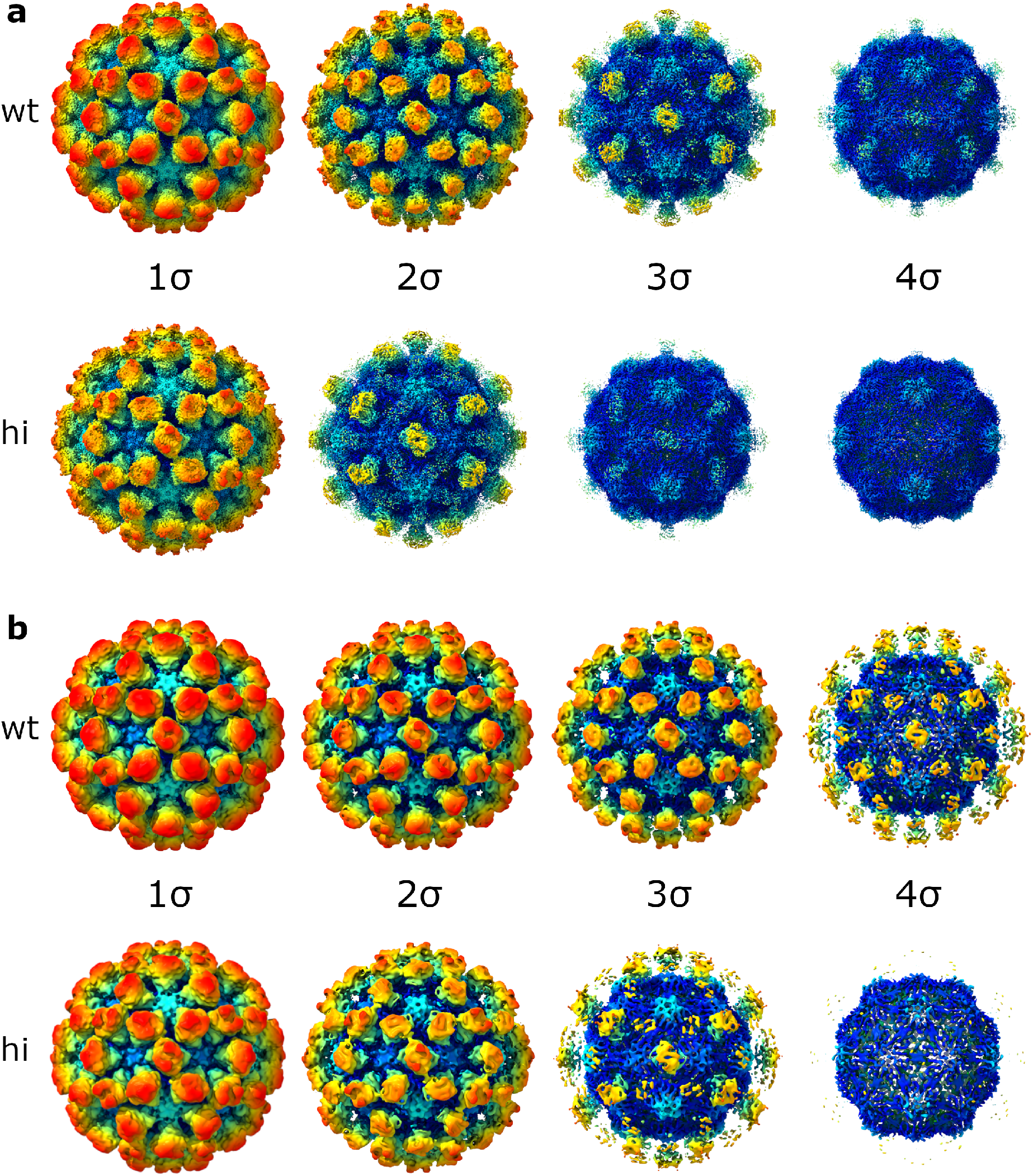
hiMNV has weaker P domain density than wtMNV. **(a)** Isosurface representations of the 3.1 Å wtMNV reconstruction (wt) and the 2.9 Å hiMNV reconstruction (hi), contoured to different thresholds (1 σ - 4 σ). **(b)** Isosurface representations of the maps shown in (a) after low-pass filtering both maps to the same resolution (8.0 Å, to allow comparison with the previously published 8.0 Å cryo-EM reconstruction of wtMNV^6^).

### Selection of a heat-stable mutant MNV

Together, our data thus far suggest P domains are highly mobile elements, however, our observations with hiMNV suggest that greater mobility is inversely correlated to infectivity. Therefore, it would follow that mutant viruses with improved thermostability would have mutations specifically affecting VP1 P domain conformation or mobility. To acquire a genetic insight into the structural determinants of P domain mobility, we generated a thermally-stabilised mutant MNV by *in vitro* evolution. A thermally-stabilised mutant MNV was isolated by repeated cycles of selection at 52°C (Figure 4a). This pool of virus, termed MNV52, showed improved thermal stability compared to wtMNV, as anticipated (Figure 4b). A PaSTRy assay revealed that viral RNA became exposed from the capsid at temperatures above 64°C (Figure 4c), consistent with the data in Figure 2b.

**Figure 4.**
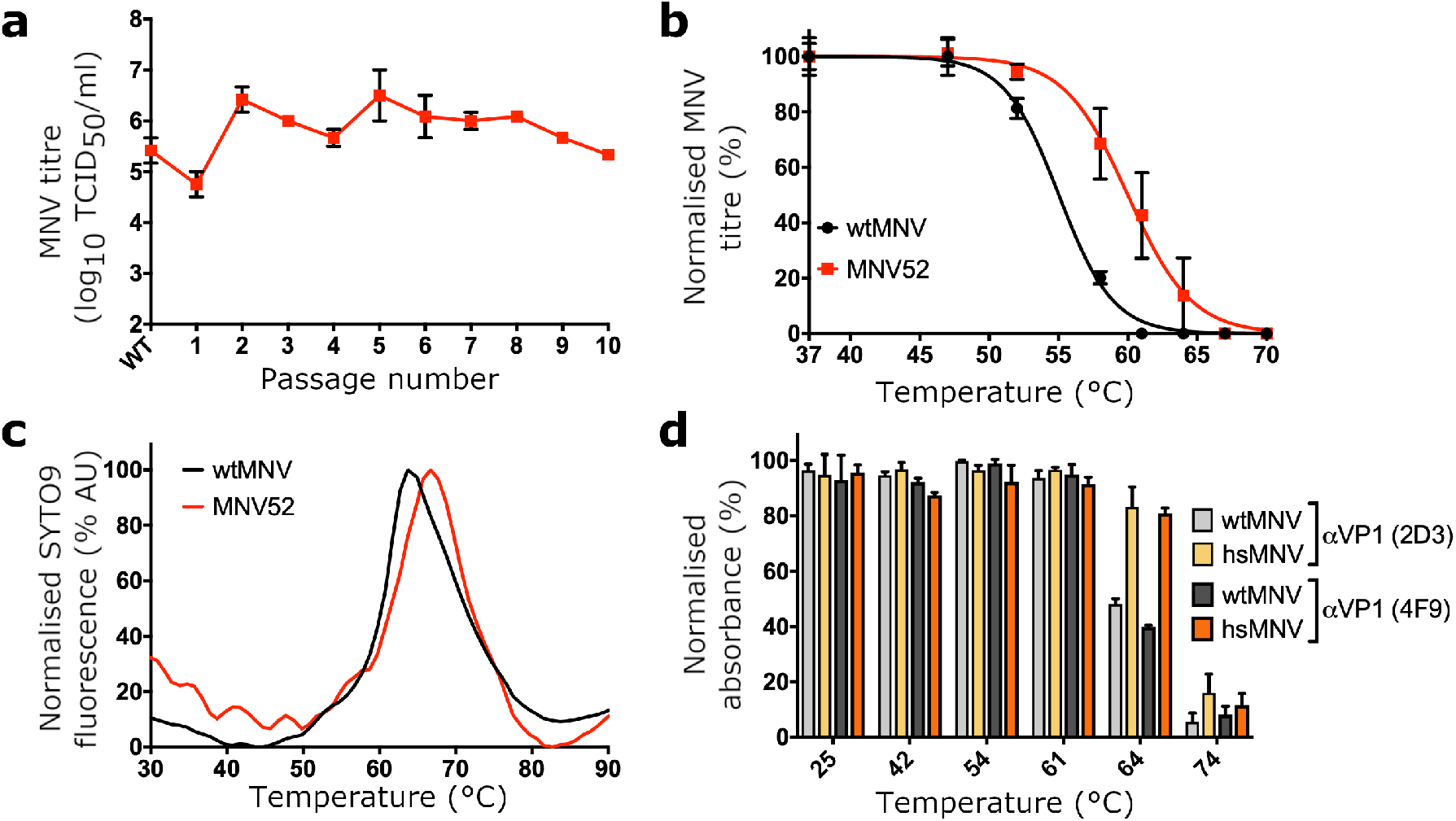
Selection and biochemical characterisation of hsMNV. **(a)** Samples of MNV were heated at 52°C for 30 minutes before cooling to 4°C. The surviving pool of viruses was subsequently passaged for 48 hours at 37°C on RAW264.7 cells. Prior to selection (‘WT’) and at each passage, the virus titre was determined by TCID_50_ on RAW264.7 cells (*n*=2 ± SEM). Consecutive cycles of selection were performed. **(b)** The pool of virus heated at 52°C between passages (MNV52) and wtMNV were heated to a range of temperatures between 37°C and 70°C and virus titre determined by TCID_50_ assay on RAW264.7 cells (*n*=4 ± SEM). **(c)** Virus samples were purified by sucrose density gradient, dialysed into PBS and used for PaSTRy thermal stability assays using the nucleic acid dye SYTO-9. **(d)** The antigenicity of wtMNV or hsMNV was determined by ELISA with anti-VP1 antibodies, 2D3 and 4F9, after incubation at the indicated temperature (*n*=2 ± SD).

To identify mutation(s) present in the thermostable MNV population, the structural protein-encoding region of the MNV genome (ORF2 and ORF3) was amplified by RT-PCR and sequenced at the consensus level. No mutation was seen in ORF3 (encoding VP2), but a single mutation was found in ORF2 that leads to a single amino acid substitution in VP1, L412Q. Consistent with our hypothesis, this mutation was located in the VP1 P domain, on the hinge loop connecting the P1 and P2 subdomains (for reference, see Figure 1i). To further characterise the effect of the L412Q substitution, we reconstituted the mutation in an infectious clone of MNV and used it to recover ‘heat-stable’ (hs)MNV particles. Like MNV52, hsMNV remained infectious after incubation at temperatures that rendered wtMNV non-infectious (Figure S5). Given that hsMNV had an amino acid substitution in the P domain of VP1, we also looked for changes in antigenicity by ELISA that may be indicative of a conformational change. We checked for binding to two neutralising anti-VP1 antibodies, 2D3 and 4F9^20^. This showed that the mutant retained two major epitopes, including at 64°C (a temperature that wtMNV could not tolerate) (Figure 4d).

### The cryo-EM reconstruction of hsMNV shows ‘twisted’ AB-type P domains

To gain a structural insight into the mechanism of stabilisation of hsMNV, we determined the structure of hsMNV to a resolution of 3.1 Å (Figure S1c, S2c). Interestingly, while CC-type P domain dimers appear virtually identical to wtMNV, AB-type P domain dimers showed a subtle difference in their orientation (Figure 5a,b, Movie S2). To explore this change, we performed rigid-body fitting of the atomic coordinates for wtMNV VP1 into the hsMNV map. This was followed by refinement against the hsMNV map with secondary structure restraints enabled. In the wtMNV map an interface is formed between A-type and C-type P domains (Figure 5c), but for hsMNV, this interface has been disrupted (Figure 5d,e). While the C-type P domain did not show any significant movement, the AB-type P domain dimer has tilted upwards, angling away from the S domains, and rotated in an anti-clockwise direction (Movie S3). As such the mutated residue now points away from the interface. In agreement with this, PDBePISA^22^ analysis of VP1 fitted into the wtMNV map suggests that C-type VP1 L412 is a buried residue and contributes to an interface with A-type VP1. When fitted into the hsMNV map, no interface is detected between A-type and C-type VP1.

**Figure 5.**
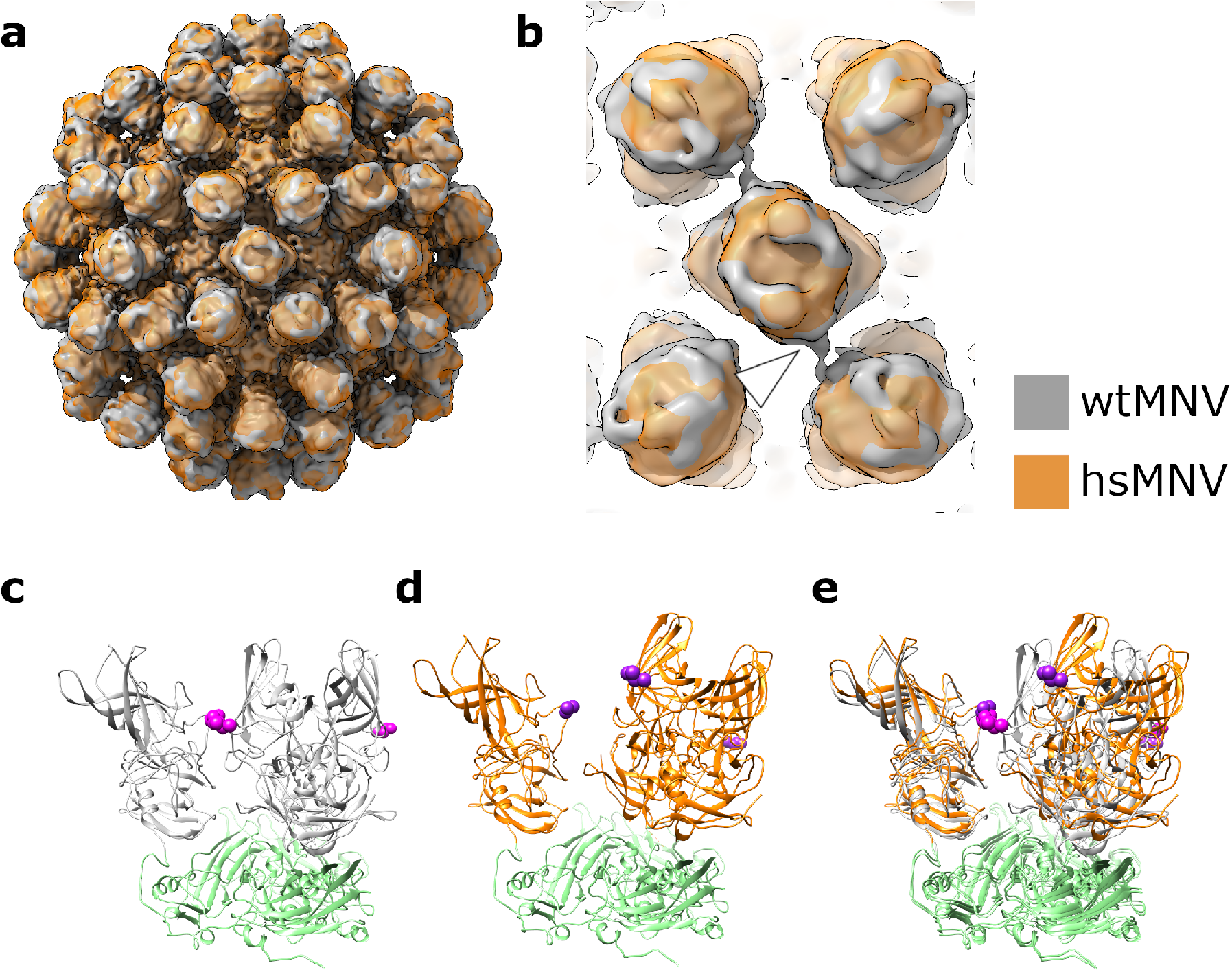
hsMNV has ‘twisted’ P domain dimers relative to wtMNV. **(a)** Overlaid isosurface representations of wtMNV (grey) and hsMNV (orange), shown at 2 σ after low-pass filtering both maps to the same resolution (8.0 Å). **(b)** Enlarged view down the icosahedral two-fold axis of the maps shown in (a), with the back focal plane clipped to remove the S domains. **(c,d)** Atomic coordinates for VP1 fitted into **(c)** wtMNV or **(d)** hsMNV density maps, centred on the AB dimer:CC dimer interface highlighted by the white arrowhead in (b). The AB-type P domain dimer and C-type P domain were fitted separately, then refined together. S domains are shown in green. The mutated residue is shown as magenta (wtMNV, L412) or dark purple (hsMNV, L412Q) spheres. **(e)** An overlay of (c) and (d).

In summary, our data suggest that the mutant virion is sampling a subset of the conformations that can be explored by the wild-type virion. Although not affecting infectivity, this is likely to impact other aspects of the viral lifecycle.

## Discussion

The structural data presented above shows the dynamic flexibility of a murine norovirus virion, by comparing three high-resolution structures of a single viral species. We showed that the infectious norovirus virion is a highly flexible macromolecular machine that is capable of sampling a range of conformational space, whilst maintaining its integrity and functionality. However, it is clear from the increased dynamics of heat-inactivated virions that too much P domain mobility is detrimental to infectivity and fitness. The virus has therefore evolved to control these dynamics and establish a balance between too much and too little P domain mobility.

MNV is a genogroup V norovirus so is closely related genetically and structurally to human noroviruses (genogroups I, II and IV), and the well-established cell culture and reverse genetic systems for MNV allowed us to specifically probe the functional significance of interactions revealed by our structural data^23,24^. Firstly, we report the high-resolution solution structure of an infectious norovirus. Our cryo-EM reconstruction of wtMNV is strikingly different to the previous, 8.0 Å resolution cryo-EM reconstruction of wtMNV, with large changes in the positioning and orientation of P domain dimers relative to the S domains^6^. This gross conformational change in P domains was consistent across all three of our independent reconstructions (discussed below) and importantly, the orientation and positioning of P domains in our wtMNV reconstruction are in line with those of most human norovirus VLPs^5,9^.

Currently, the reasons for the differences between our reconstruction and that in Katpally *et al*, 2010^6^, are unclear. The previously reported wtMNV reconstruction had a conformation similar to a genogroup (G)II.10 norovirus VLP complexed with an antigen-binding fragment (Fab) of a monoclonal antibody, and to a GII.4 Minerva VLP, though it should be noted that the latter formed a *T* = 4 capsid that is perhaps less relevant to the *T* = 3 capsid of the infectious virion^8,25^. However, our reconstruction is consistent with all other norovirus VLP structures. Given the dynamic nature of the P domains shown here, it seems plausible that the two reconstructions represent different functionally-relevant conformations that the MNV capsid can adopt. These could have been induced experimentally during handling of the virus. For example, there were small differences in virus purification protocols, and we cannot rule out that the two reconstructions were on viruses with subtly different VP1 primary sequences (e.g. arising from mutation during passage), despite both starting with a CW1 strain of MNV-1^7^. It is also possible that differences in buffer composition are responsible for the alternative conformations. Given the changing ionic composition of the environment that the virus is exposed to during endocytosis into a target cell^26^, this latter point may be of biological importance. Using the crystal structure of an MNV P domain dimer complexed with the MNV receptor, CD300lf (PDB: 6C6Q)^11^, we generated a receptor docking model based on the MNV reconstruction reported here (Figure S6). Interestingly, a clash between trimers of CD300lf in the previous docking model^11^ is resolved in this model, and there is space for higher receptor occupancy.

The independent mobility of individual P domain dimers on the viral capsid surface is striking. While it is suspected that individual norovirus species can adopt multiple different gross morphologies^15^, here we provide direct structural evidence for alternative morphologies within a single norovirus species, and within individual norovirus particles. Using focussed 3D classification, we identified a remarkable diversity in P domain dimer positioning, which was not co-ordinated over the capsid surface, and was not equal between different quasi-conformers, with CC-type dimers less mobile than AB-type dimers. This contrasts with data obtained for another calicivirus, Tulane virus, where CC-type dimers were shown to be more mobile^27^. Surprisingly, one class (out of ten) for each of the AB- and CC-type dimer classifications had an inverted orientation in the *z*-axis (Figure S3a,b). However, for the remaining classes, no significant improvement in P domain density was observed – likely because any improvement from removing heterogeneity in the reconstruction is countered by a decreased number of particles contributing to the final reconstruction.

By determining the structure of the wtMNV capsid using cryo-EM, we were able to identify unambiguous differences between quasi-equivalent subunits of VP1. The N-terminal regions of VP1 were particularly variable – in A-type VP1, the N-terminal arm protrudes deeper into the capsid to interact with other A-type subunits around the icosahedral five-fold axis. B-type VP1 N-terminal regions run towards the icosahedral three-fold axis, and C-type VP1 N-terminal regions are not resolved in our map, suggesting increased flexibility. We note that this difference between quasi-equivalent subunits could provide a mechanism to guide the positioning of other components of the virion, such as VP2. Based on work with feline calicivirus, it has been proposed that VP2 may bind the capsid interior at a single icosahedral three-fold axis^28^. While it has been shown through mutational analysis that VP1 N-termini are not required for co-precipitation with VP2^13^, the ability of VP1 N-termini to organise differently may provide an interface for ‘recognition’ of the three-fold axis (formed by B- and C-type VP1 subunits); i.e., while not required for VP2 packaging, one may speculate that differences between N-terminal regions of VP1 could guide the positioning of VP2 within the capsid interior. Alternatively, these differences may guide the positioning of VPg or the RNA genome, which may in turn guide VP2.

To support our hypothesis that viral P domains are mobile elements that can adopt multiple conformations, we endeavoured to induce conformational changes reflective of those that may occur during infection by thermally stressing MNV, leading to the generation of non-infectious virus with intact capsids (hiMNV). While cryo-EM revealed no morphological differences, P domain density was weaker than for wtMNV. Weaker density may arise from ‘partial occupancy’ (i.e. inclusion of particles that have lost some or all P domains in the final reconstruction), or more likely from increased P domain flexibility/mobility. While we saw small particles reminiscent of capsid proteins in the background of raw micrographs from the hiMNV data set (Figure S1b), we did not observe any P domain-lacking capsids in raw micrographs and failed to pull out classes lacking P domains in asymmetric 3D classification, so believe the latter explanation to be the most plausible. It is not immediately clear why increased P domain mobility would render the virus non-infectious – perhaps the range of conformations sampled by the P domains is changed, such that a conformation required for infection is no longer accessible. However, we cannot rule out the possibility of other subtle structural changes within the unresolved regions of P domains that contribute to virus inactivation. These potential changes may disrupt the interaction between the virus and its receptor, or could cause a change in the amount/composition of metal ions coordinated by VP1, which are important for viral infectivity and capsid stability^8,11^. We also cannot discount the possibility that structural changes to other capsid components/contents (such as VP2 or VPg which are not resolved in our icosahedrally-averaged reconstructions) may contribute to the loss of infectivity, though we believe that changes to VP1 are ultimately responsible.

In support of this hypothesis, MNV passaged with a thermal selection pressure acquired resistance to heat inactivation from a single point mutation in the gene encoding VP1 (ORF2). Our cryo-EM reconstruction of hsMNV showed that AB-type P domain dimers were ‘twisted/tilted’ relative to their positions in wtMNV, disrupting an interface between AB- and CC-type dimers. While it is theoretically possible that this altered P domain dimer positioning is unrelated to the mutation, the most plausible interpretation is that the L412Q substitution (which is located within this interface) is responsible. It is not obvious why the mutant morphology is more stable – one would expect disruption of an interface to decrease stability. Indeed, thermostabilised foot-and-mouth disease virus (FMDV) capsids were generated by introducing a disulphide bond to stabilise an interface between adjacent pentamers^29^, and many stabilising mutations seen in poliovirus are thought to act by stabilising interfaces between subunits^30^. We suspect that the ‘twisted’ morphology of hsMNV liberates P domain dimers to enter a protective conformation upon heating, that may only become apparent at high temperatures. Alternatively, there may be subtle, protective conformational changes within regions of the P domains that are unresolved in our map. Another possibility is that the mutant morphology may facilitate new stabilising interactions with other viral components, such as the portal thought to be formed by VP2 during genome release^14^. It should be noted that we do not detect VP2 in any of the structures reported here.

Here, we were interested in a thermostabilised MNV variant for the insights it provides about norovirus structural biology. However, it also offers a platform for the development of a thermostabilised vaccine against noroviruses. Most norovirus vaccine candidates currently under development are based on VLPs^31^. VLPs lack a viral genome, which can provide stabilising interactions with the capsid^32^, potentially resulting in reduced capsid stability. This may be a problem during vaccine distribution. Norovirus vaccines are most urgently needed in countries with warmer climates and require a cold chain. This can be particularly challenging to maintain in hard-to-reach regions, so a thermally stabilised vaccine is a particularly attractive prospect. By incorporating the L412Q mutation into VLP production, it may be possible to generate more stable VLPs. Essential for this strategy, mutant VLPs must generate an appropriate neutralising antibody response. In this regard, hsMNV retained its major antigenic determinants when tested against two neutralising antibodies, which is important for future vaccine production. Furthermore, VP1 sequences from human noroviruses (genogroups I, II and IV) show conservation of a hydrophobic residue in the same position (Figure S7). Thus, it is plausible that the same mutation may stabilise human norovirus VLPs for use in a stabilised human norovirus VLP-based vaccine.

Overall, this work shows that norovirus P domains appear to be dynamic components of the capsid, continually sampling different conformations and positions relative to the S domains. This may benefit the virus in a number of ways, for example, by contributing to immune evasion and facilitating binding to the target cell, as outlined in a recent review article^15^. In addition to providing insights into norovirus biology, our structures and description of a stabilising VP1 mutation will prove useful resources for the future study of noroviruses. Furthermore, we propose that incorporation of such stabilising mutations could compensate for the lack of genome in VLPs, generating a more stable particle that could pave the way for the development of vaccines against norovirus disease.

## Methods

### Cell lines and antibodies

RAW264.7 cells (kindly gifted by Ian Clarke, University of Southampton) and BHK-21 cells (obtained from the ATCC) were maintained in high-glucose Dulbecco’s Modified Eagle’s Medium (DMEM) supplemented with 10% (v/v) fetal bovine serum, 20 mM HEPES buffer and 50 U/ml penicillin and streptomycin. Cells were incubated at 37°C, 5% CO_2_. Neutralising antibodies against MNV VP1, 2D3 and 4F9^20^, were kindly gifted by Christiane Wobus (University of Michigan).

### MNV propagation

To generate virus for use in this study, MNV-1 strain CW1P3^33^ (referred to as MNV) was recovered from an infectious clone, and propagated in RAW264.7 cells as described previously^23^. Briefly, RAW264.7 cells were seeded in T175 flasks and allowed to reach 80% confluency. They were then infected with crude stocks of MNV in fresh media and incubated for 48 - 72 hours, then harvested when confluent cytopathic effect was visible. The infectious media was put through three freeze-thaw cycles to release virus. To concentrate virus for propagation at high multiplicity of infection (MOI), the infectious lysate was first clarified (3300 × *g*, 10 min, 4°C), then taken for ultracentrifugation at 366,000 × *g* (60 min, 4°C) and the pellet resuspended in phosphate-buffered saline (PBS).

To minimise the chance of reversion, concentrated crude stocks of hsMNV were heated to 52°C for 30 min between each passage. Both wtMNV and hsMNV were validated by sequencing prior to structural analysis.

### MNV purification

To purify MNV, we followed modified versions of the protocols described by Hwang *et al.*^34^. Infectious media was collected and freeze-thawed three times to lyse cells and release virus. NP-40 was added to the infectious lysate to a final concentration of 0.1%, before three rounds of centrifugation (3300 × *g*, 10 min each, 4°C), each time discarding the pelleted cell debris. The clarified supernatant was loaded onto a 30% (w/v) sucrose cushion and subjected to ultracentrifugation at 150 000 × *g* (3 h, 4°C). The resultant pellets were resuspended in PBS, then clarified by centrifugation at 17,000 × *g* (10 min), before being loaded onto a continuous 15 - 60% sucrose gradient for ultracentrifugation at 300,000 × *g* (50 min, 4°C) then fractionated. Peak fractions (determined by SDS-PAGE analysis) were combined, spun at 366,000 × *g* (60 min, 4°C) and the pellet resuspended in PBS for a second round of sucrose gradient ultracentrifugation.

To remove sucrose for structural studies, peak fractions from the second sucrose gradient were combined and dialysed using a 10,000 molecular weight cut-off Slide-A-Lyzer dialysis cassette (Thermo Fisher) in 1 litre EM buffer (10 mM HEPES [pH 7.6], 200 mM NaCl, 5 mM MgCl_2_, 1 mM KCl, 1 mM CaCl_2_). After 1 hour at room temperature, the cassette was transferred into fresh EM buffer for another hour at room temperature, before being transferred to fresh EM buffer for overnight incubation at 4°C. Virus was recovered from the dialysis cassette and stored at 4°C prior to imaging.

### Median tissue culture infectious dose (TCID_50_) assay

To measure viral infectivity, TCID_50_ assays were performed, according to a modified version of the protocol described by Hwang *et al.*^34^. RAW264.7 cells were seeded into 96-well plates at a density of 2.0 × 10^4^ cells/well and incubated for 24 h. Subsequently, 10-fold serial dilutions of MNV in fresh DMEM were prepared, and 100 μl of each concentration was added to 100 μl of media already present in each well. For each concentration of MNV, six wells were infected. Cells were incubated for a further 72 h before fixing with 4% paraformaldehyde (PFA) in PBS, and staining with crystal violet solution to assess cytopathic effect. TCID_50_ values were calculated according to the Spearman & Kärber algorithm^35^.

### Particle Stability Thermal Release (PaSTRy) assay

To investigate particle stability, PaSTRy assays^19^ were performed according to the protocol described by Adeyemi *et al.*^36^. Briefly, 1.0 μg purified MNV was incubated in a 50 μl reaction mixture of 5 μM SYTO9, 150× SYPRO-Orange and PaSTRy buffer (2 mM HEPES [pH 8.0], 200 mM NaCl) on a temperature ramp from 25°C to 95°C. At every 1°C interval, the Stratagene MX3005p quantitative-PCR (qPCR) system was used to measure fluorescence.

### RNase protection assay

To investigate the disassembly of virus particles, an RNase protection assay was performed based on the protocol reported by Groppelli *et al.*^37^, with several modifications. Briefly, purified MNV was heated to a range of temperatures. After heating, a fraction of each sample was retained for titration by TCID_50_ assay (‘Virus Infectivity’), while the remaining fraction was treated with RNase A (1 mg/ml) for 30 min at 37°C. To stop the reaction and extract viral RNA, TRIzol was added and RNA extracted using the Direct-zol RNA miniprep kit (Zymo Research) according to the manufacturer’s instructions. Total extracted RNA was transfected into BHK-21 cells using lipofectin (as described previously^38^), supplemented with carrier RNA (yeast tRNA) to 1 μg per reaction. 48 hours later, total virus was extracted by freeze-thaw, cell debris clarified by centrifugation, and virus titrated by TCID_50_ assay.

### Selection of thermally-stable MNV (hsMNV)

To generate a thermally-stable population of MNV, crude MNV samples were heated at 52°C for 30 minutes before cooling to 4°C. The surviving pool of virus was subsequently passaged at 37°C on RAW264.7 cells. Consecutive cycles of selection and passage were performed, after which the pool of virus was characterised.

### Enzyme-Linked Immunosorbent Assays (ELISAs)

To check for changes to antigenicity, ELISAs were performed according to the protocol described by Hwang *et al.*^34^. Concentrated MNV suspended in PBS was used to coat ELISA wells overnight at 4°C, which were then washed with ELISA wash buffer (150 mM NaCl, 0.05% Tween 20) and blocked with ELISA blocking buffer (50 mM Na_2_CO_3_, 50 mM NaHCO_3_, 3% BSA, pH 11) at 37°C for 2 hours. After blocking, wells were washed twice with ELISA wash buffer, then incubated with primary antibody (2D3 or 4F9) diluted 1:100 in ELISA III buffer (150 mM NaCl, 1 mM EDTA, 50 mM Tris-HCl, 0.05% Tween 20, 0.1% BSA, pH 11) for 1 hour at 37°C. Primary antibody was removed and wells were washed four times with ELISA wash buffer, before adding secondary antibody (peroxidase-conjugated anti-mouse IgG [A9044, Sigma-Aldrich]) diluted 1:2000 in ELISA III buffer for 1 hour at 37°C. Secondary antibody was removed and wells were washed four times with ELISA wash buffer, then substrate (ABTS) was added and incubated at room temperature for 30-60 min. Reactions were stopped with 0.2 N phosphoric acid and absorbance in each well measured by plate reader at 415 nm.

### Cryo-electron microscopy

To prepare MNV samples for cryo-EM, lacey carbon 400-mesh copper grids coated with a <3 nm continuous carbon film (Agar Scientific, UK) were glow-discharged in air or amylamine vapour (10 mA, 30 seconds) before applying two to three 3 μl aliquots of purified MNV, to improve the concentration of virus on the grid surface (as described previously^39^). Each application was followed by a 30 second incubation period at 80% relative humidity (8°C), then the grid was manually blotted to remove excess fluid before the next application. 30 seconds after the final application, grids were blotted and vitrified in liquid nitrogen-cooled liquid ethane using a LEICA EM GP plunge freezing device (Leica Microsystems). Grids were stored in liquid nitrogen prior to imaging with an FEI Titan Krios transmission electron microscope (ABSL, University of Leeds) at 300 kV, at a magnification of 75 000× and a calibrated object sampling of 1.065 Å/pixel. A complete set of data collection parameters for each sample is provided in Table S1.

### Image processing

Following cryo-EM data collection, the RELION-2.1 and RELION-3.0 pipelines^40–42^ were used for image processing. Drift correction was first performed on micrograph stacks using MOTIONCOR2^43^, and the contrast transfer function for each was estimated using Gctf^44^. A subset of virus particles was picked manually and subject to 2D classification, with resultant classes used as templates for automatic particle picking^45^. Particles were classified through multiple rounds of reference-free 2D classification, and particles in poor quality classes were removed after each round. An initial 3D model was generated *de novo*^46^ and used as a reference for 3D auto-refinement with icosahedral symmetry imposed. This reconstruction was post-processed to mask and correct for the B-factor of the map, before (i) taking particles forward to CTF refinement and Bayesian polishing, or (ii) further ‘clean-up’ by alignment-free 3D classification, with particles from subsequent 3D auto-refinement and post-processing being used for CTF refinement and Bayesian polishing. Multiple rounds of CTF refinement (with or without beamtilt refinement) and Bayesian polishing^47^ were performed, before final icosahedral symmetry-imposed 3D auto-refinement and post-processing. The nominal resolution for each map was determined according to the ‘gold standard’ Fourier shell correlation criterion (FSC = 0.143)^48^, and the local resolution estimation tool in RELION was used to generate maps filtered by local resolution.

To investigate P domain mobility, a focussed 3D classification approach was employed (as described previously^14,49–51^). Briefly, each particle contributing to the final icosahedral symmetry-imposed reconstruction was assigned 60 orientations corresponding to its icosahedrally-related views using the relion_symmetry_expand tool. SPIDER^52^ was used to generate a cylindrical mask to isolate either an AB-type or a CC-type P domain dimer, and the symmetry expanded particles were subjected to masked 3D classification without alignment, using a regularisation parameter (‘T’ number) of 20. Classes were inspected visually, and particles from selected classes (with assigned orientation information) were used to generate full capsid reconstructions without imposing symmetry, using the relion_reconstruct tool.

### Model building and refinement

To generate a preliminary model for the VP1 asymmetric unit, the amino acid sequence corresponding to the S domain of MNV VP1 was used to build a homology model with the Phyre2 server^16^, which was rigid-body fitted into each quasi-equivalent position in the wtMNV density map using UCSF Chimera^53^. This preliminary model was manually refined in Coot^54^, symmetrised in UCSF Chimera to generate the other 59 copies of the asymmetric unit that form the capsid, then subject to ‘real space refinement’ in Phenix^55^. To improve the geometry of the coordinates and fit of the model to the density map, the S domain model was iterated between manual fitting in Coot and refinement in Phenix. Following this, the crystal structure of an MNV P domain complexed with CD300lf (PDB: 6C6Q)^11^ was also fitted into the map to occupy each quasi-equivalent position of the asymmetric unit, after removing ligands/CD300lf and correcting the peptide sequence. P domain coordinates were combined with the refined S domain model, and subject to a single round of refinement in Phenix. For each real space refinement, secondary structure restraints were imposed. Molprobity^56^ was used to validate the model.

### Analysis and visualisation

For structural analysis and generation of figures, density maps and atomic coordinates were viewed in UCSF Chimera^53^, UCSF ChimeraX^57^ and PyMOL (The PyMOL Molecular Graphics System, Version 2.0, Schrödinger, LLC). RMSD values between quasi-equivalent states were calculated using the ‘MatchMaker’ tool of UCSF Chimera with default settings.

### Structure deposition

Cryo-EM density maps for wtMNV, hiMNV and hsMNV will be deposited to the Electron Microscopy Data Bank (EMDB), and atomic coordinates for wtMNV VP1 will be deposited to the Protein Data Bank (PDB).

## Acknowledgements

This work was supported by Wellcome Trust PhD studentships to JSS (102174/B/13/Z) and DLH (102572/B/13/Z), and MRC funding to MRH (MR/S007229/1). Electron microscopy was performed in the Astbury Biostructure Laboratory, which was funded by the University of Leeds and the Wellcome Trust (108466/Z/15/Z).

## Author contributions

JSS, DLH, OOA, NAR, MRH and NJS designed the study and wrote the manuscript. MRH and OOA generated hsMNV and performed thermostability assays. JSS, MRH and OOA propagated and purified virus for structural studies. JSS and DLH collected cryo-EM data, carried out image processing and atomic modelling, and analysed the structures. NAR, MRH and NJS provided supervision.

## Materials & correspondence

Correspondence and materials requests should be directed to MRH and NJS.

## References

1. Patel, M. M. et al. Systematic literature review of role of noroviruses in sporadic gastroenteritis. Emerg. Infect. Dis. 14, 1224–31 (2008).

2. Atmar, R. L. et al. Norovirus Vaccine against Experimental Human Norwalk Virus Illness. N. Engl. J. Med. 365, 2178–87 (2011).

3. Bernstein, D. I. et al. Norovirus Vaccine Against Experimental Human GII.4 Virus Illness: A Challenge Study in Healthy Adults. J. Infect. Dis. 211, 870–8 (2015).

4. Lambden, P. R., Caul, E. O., Ashley, C. R. & Clarke, I. N. Sequence and genome organization of a human small round-structured (Norwalk-like) virus. Science 259, 516–9 (1993).

5. Prasad, B. V., Rothnagel, R., Jiang, X. & Estes, M. K. Three-dimensional structure of baculovirus-expressed Norwalk virus capsids. J. Virol. 68, 5117–25 (1994).

6. Katpally, U. et al. High-Resolution Cryo-Electron Microscopy Structures of Murine Norovirus 1 and Rabbit Hemorrhagic Disease Virus Reveal Marked Flexibility in the Receptor Binding Domains. J. Virol. 84, 5836–41 (2010).

7. Katpally, U., Wobus, C. E., Dryden, K., Virgin, H. W. & Smith, T. J. Structure of Antibody-Neutralized Murine Norovirus and Unexpected Differences from Viruslike Particles. J. Virol. 82, 2079–88 (2008).

8. Jung, J. et al. High-resolution cryo-EM structures of outbreak strain human norovirus shells reveal size variations. Proc. Natl. Acad. Sci. USA 116, 12828–32 (2019).

9. Prasad, B. V. et al. X-ray Crystallographic Structure of the Norwalk Virus Capsid. Science 286, 287–90 (1999).

10. Kilic, T., Koromyslova, A. & Hansman, G. S. Structural basis for human norovirus capsid binding to bile acids. J. Virol. 93, e01581–18 (2019).

11. Nelson, C. A. et al. Structural basis for murine norovirus engagement of bile acids and the CD300lf receptor. Proc. Natl. Acad. Sci. USA 115, E9201–10 (2018).

12. Koromyslova, A. D., Leuthold, M. M., Bowler, M. W. & Hansman, G. S. The sweet quartet: Binding of fucose to the norovirus capsid. Virology 483, 203–8 (2015).

13. Vongpunsawad, S., Venkataram Prasad, B. V. & Estes, M. K. Norwalk Virus Minor Capsid Protein VP2 Associates within the VP1 Shell Domain. J. Virol. 87, 4818–25 (2013).

14. Conley, M. J. et al. Calicivirus VP2 forms a portal-like assembly following receptor engagement. Nature 565, 377–381 (2019).

15. Smith, H. Q. & Smith, T. J. The Dynamic Capsid Structures of the Noroviruses. Viruses 11, E235 (2019).

16. Kelley, L. A., Mezulis, S., Yates, C. M., Wass, M. N. & Sternberg, M. J. The Phyre2 web portal for protein modeling, prediction and analysis. Nat. Protoc. 10, 845–58 (2015).

17. Curry, S., Chow, M. & Hogle, J. M. The poliovirus 135S particle is infectious. J. Virol. 70, 7125–31 (1996).

18. Bubeck, D. et al. The structure of the poliovirus 135S cell entry intermediate at 10-angstrom resolution reveals the location of an externalized polypeptide that binds to membranes. J. Virol. 79, 7745–55 (2005).

19. Walter, T. S. et al. A plate-based high-throughput assay for virus stability and vaccine formulation. J. Virol. Methods 185, 166–70 (2012).

20. Kolawole, A. O. et al. Newly isolated mAbs broaden the neutralizing epitope in murine norovirus. J. Gen. Virol. 95, 1958–68 (2014).

21. Taube, S. et al. High-Resolution X-Ray Structure and Functional Analysis of the Murine Norovirus 1 Capsid Protein Protruding Domain. J. Virol. 84, 5695–705 (2010).

22. Krissinel, E. & Henrick, K. Inference of Macromolecular Assemblies from Crystalline State. J. Mol. Biol. 372, 774–97 (2007).

23. Wobus, C. E. et al. Replication of Norovirus in cell culture reveals a tropism for dendritic cells and macrophages. PLoS Biol. 2, e432 (2004).

24. Chaudhry, Y., Skinner, M. A. & Goodfellow, I. G. Recovery of genetically defined murine norovirus in tissue culture by using a fowlpox virus expressing T7 RNA polymerase. J. Gen. Virol. 88, 2091–100 (2007).

25. Hansman, G. S. et al. Structural Basis for Broad Detection of Genogroup II Noroviruses by a Monoclonal Antibody That Binds to a Site Occluded in the Viral Particle. J. Virol. 86, 3635–46 (2012).

26. Scott, C. C. & Gruenberg, J. Ion flux and the function of endosomes and lysosomes: pH is just the start: the flux of ions across endosomal membranes influences endosome function not only through regulation of the luminal pH. BioEssays 33, 103–10 (2011).

27. Yu, G. et al. Cryo-EM Structure of a Novel Calicivirus, Tulane Virus. PLoS One 8, e59817 (2013).

28. Conley, M. et al. Calicivirus VP2 forms a portal to mediate endosome escape. Preprint at bioRxiv. doi:10.1101/397901

29. Porta, C. et al. Rational Engineering of Recombinant Picornavirus Capsids to Produce Safe, Protective Vaccine Antigen. PLoS Pathog. 9, e1003255 (2013).

30. Fox, H., Knowlson, S., Minor, P. D. & Macadam, A. J. Genetically Thermo-Stabilised, Immunogenic Poliovirus Empty Capsids; a Strategy for Non-replicating Vaccines. PLoS Pathog. 13, e1006117 (2017).

31. Mattison, C. P., Cardemil, C. V. & Hall, A. J. Progress on norovirus vaccine research: public health considerations and future directions. Expert Rev. Vaccines 17, 773–84 (2018).

32. Snijder, J. et al. Probing the biophysical interplay between a viral genome and its capsid. Nat. Chem. 5, 502–9 (2013).

33. Ward, V. K. et al. Recovery of infectious murine norovirus using pol II-driven expression of full-length cDNA. Proc. Natl. Acad. Sci. USA 104, 11050–5 (2007).

34. Hwang, S. et al. Murine Norovirus: Propagation, Quantification, and Genetic Manipulation. Curr. Protoc. Microbiol. 33, 15K.2.1–61 (2014).

35. Hierholzer, J. C. & Killington, R. A. 2 - Virus isolation and quantitation. in Virology Methods Manual 25–46 (Academic Press, 1996).

36. Adeyemi, O. O., Nicol, C., Stonehouse, N. J. & Rowlands, D. J. Increasing Type 1 Poliovirus Capsid Stability by Thermal Selection. J. Virol. 91, e01586–16 (2017).

37. Groppelli, E. et al. Picornavirus RNA is protected from cleavage by ribonuclease during virion uncoating and transfer across cellular and model membranes. PLoS Pathog. 13, e1006197 (2017).

38. Herod, M. R. et al. Genetic economy in picornaviruses: Foot-and-mouth disease virus replication exploits alternative precursor cleavage pathways. PLoS Pathog. 13, e1006666 (2017).

39. Hurdiss, D. L., Frank, M., Snowden, J. S., Macdonald, A. & Ranson, N. A. The Structure of an Infectious Human Polyomavirus and Its Interactions with Cellular Receptors. Structure 26, 839–47.e3 (2018).

40. Kimanius, D., Forsberg, B. O., Scheres, S. H. W. & Lindahl, E. Accelerated cryo-EM structure determination with parallelisation using GPUs in RELION-2. Elife 5, e18722 (2016).

41. Scheres, S. H. W. RELION: Implementation of a Bayesian approach to cryo-EM structure determination. J. Struct. Biol. 180, 519–30 (2012).

42. Zivanov, J. et al. New tools for automated high-resolution cryo-EM structure determination in RELION-3. Elife 7, e42166 (2018).

43. Zheng, S. Q. et al. MotionCor2: anisotropic correction of beam-induced motion for improved cryo-electron microscopy. Nat. Methods 14, 331–2 (2017).

44. Zhang, K. Gctf: Real-time CTF determination and correction. J. Struct. Biol. 193, 1–12 (2016).

45. Scheres, S. H. W. Semi-automated selection of cryo-EM particles in RELION-1.3. J. Struct. Biol. 189, 114–22 (2015).

46. Punjani, A., Rubinstein, J. L., Fleet, D. J. & Brubaker, M. A. cryoSPARC: algorithms for rapid unsupervised cryo-EM structure determination. Nat. Methods 14, 290–96 (2017).

47. Zivanov, J., Nakane, T. & Scheres, S. H. W. A Bayesian approach to beam-induced motion correction in cryo-EM single-particle analysis. IUCrJ 6, 5–17 (2019).

48. Scheres, S. H. W. & Chen, S. Prevention of overfitting in cryo-EM structure determination. Nat. Methods 9, 853–4 (2012).

49. McElwee, M., Vijayakrishnan, S., Rixon, F. & Bhella, D. Structure of the herpes simplex virus portal-vertex. PLoS Biol. 16, e2006191 (2018).

50. Scheres, S. H. W. Processing of Structurally Heterogeneous Cryo-EM Data in RELION. Methods Enzymol. 579, 125–57 (2016).

51. Zhou, M. et al. Atomic structure of the apoptosome: mechanism of cytochrome c- and dATP-mediated activation of Apaf-1. Genes Dev. 29, 2349–61 (2015).

52. Frank, J. et al. SPIDER and WEB: Processing and Visualization of Images in 3D Electron Microscopy and Related Fields. J. Struct. Biol. 116, 190–9 (1996).

53. Pettersen, E. F. et al. UCSF Chimera - A visualization system for exploratory research and analysis. J. Comput. Chem. 25, 1605–12 (2004).

54. Emsley, P., Lohkamp, B., Scott, W. G. & Cowtan, K. Features and development of Coot. Acta Crystallogr. D. Biol. Crystallogr. 66, 486–501 (2010).

55. Adams, P. D. et al. PHENIX: a comprehensive Python-based system for macromolecular structure solution. Acta Crystallogr. D. Biol. Crystallogr. 66, 213–21 (2010).

56. Chen, V. B. et al. MolProbity: all-atom structure validation for macromolecular crystallography. Acta Crystallogr. D. Biol. Crystallogr. 66, 12–21 (2010).

57. Goddard, T. D. et al. UCSF ChimeraX: Meeting modern challenges in visualization and analysis. Protein Sci. 27, 14–25 (2018).

