## Supplementary Material for "High-Resolution Cryo-EM Reveals Dynamics in the Murine Norovirus Capsid"

*Joseph S. Snowden, Daniel L. Hurdiss, Oluwapelumi O. Adeyemi, Neil A. Ranson, Morgan R. Herod & Nicola J. Stonehouse*

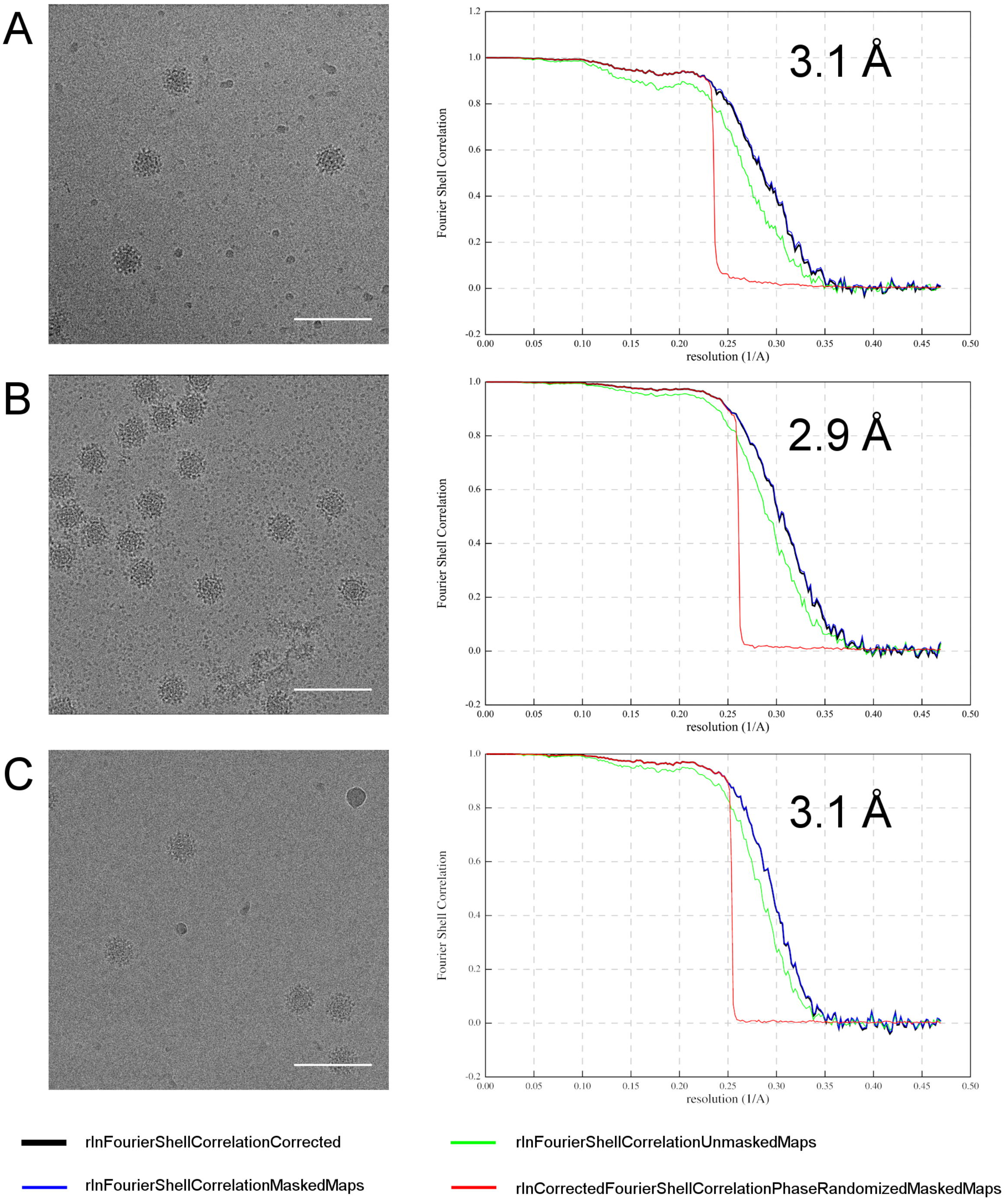


**Figure S1.** Representative micrographs and Fourier shell correlation (FSC) plots corresponding to data sets for wtMNV **(A)**, hiMNV **(B)**, and hsMNV **(C)**. Scale bars show 100 nm. The resolution given for each data set is determined using the FSC=0.143 criterion.

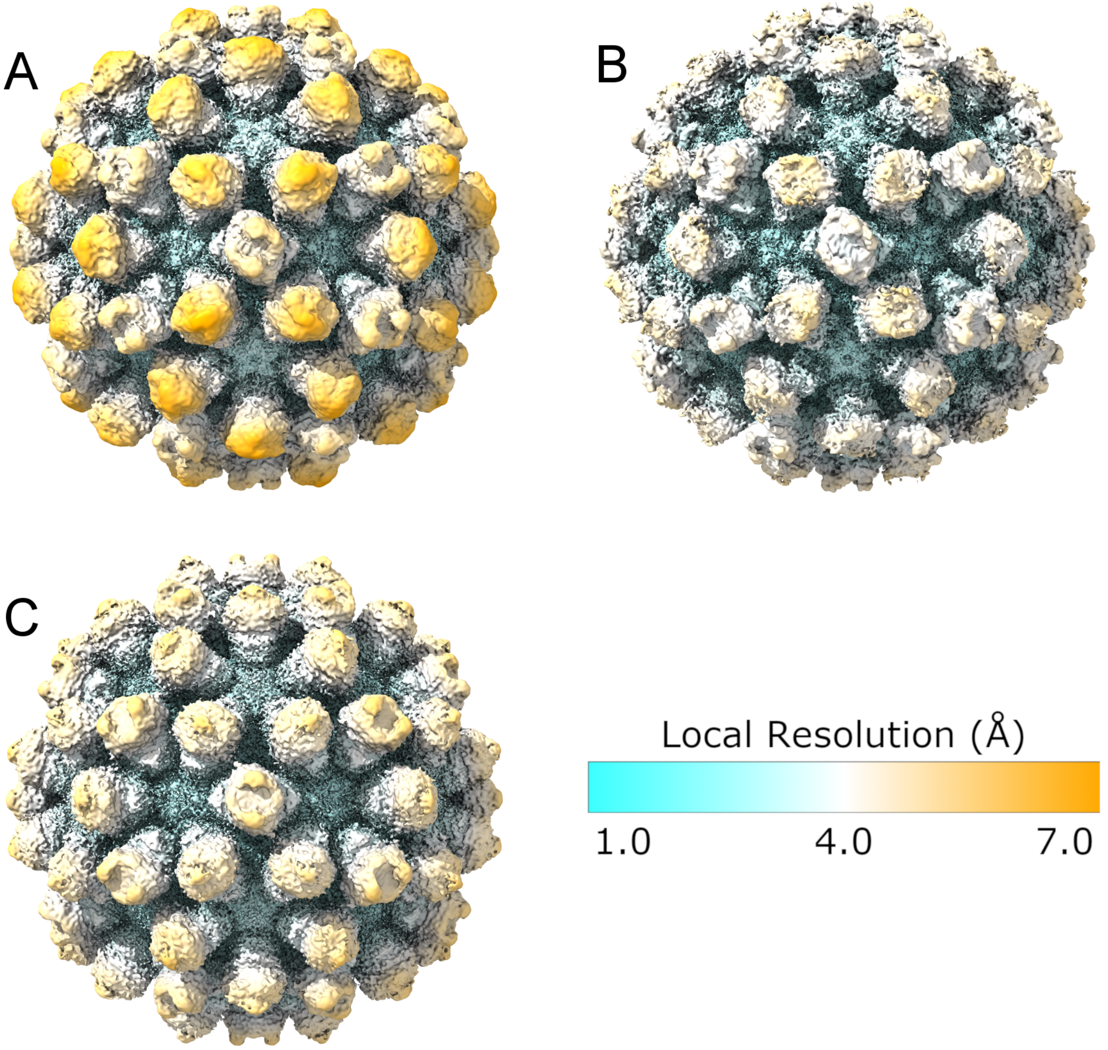


**Figure S2.** Isosurface representations of **(A)** the 3.1 Å wtMNV reconstruction, **(B)** the 2.9 Å hiMNV reconstruction and **(C)** 3.1 Å hsMNV reconstruction, coloured according to local resolution. All reconstructions are shown at 1 $\sigma$.

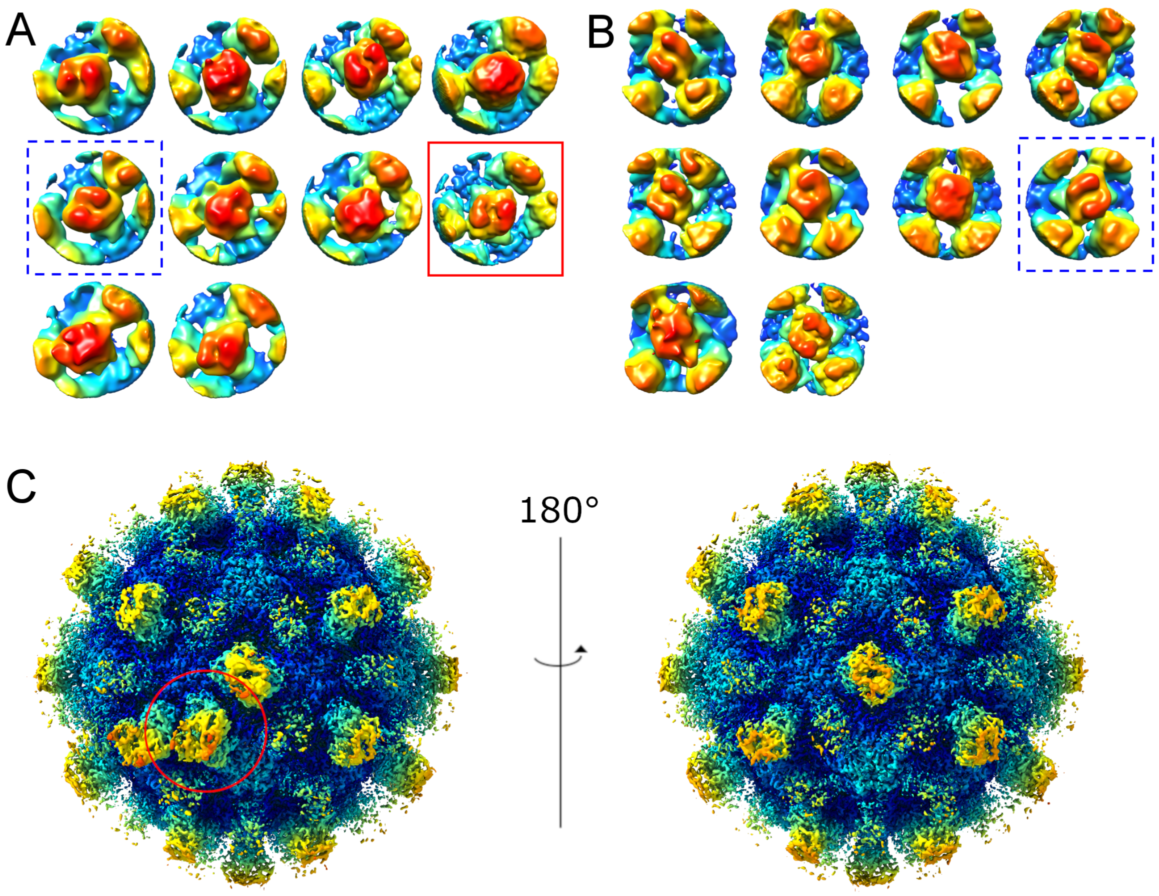


**Figure S3. Focussed classification of wtMNV. (A,B)** All classes resulting from focussed classification of (A) AB P domain dimers and (B) CC P domain dimers from wtMNV, shown at 5 $\sigma$ and coloured according to height. Classes with an inverted *Z* orientation are indicated by a dashed blue box. **(C)** Reconstructed capsid from a single AB P domain dimer class (shown by the red box in (A)) shown at 2.8 $\sigma$ with a radial colour scheme. The area used for focussed classification is indicated by a red circle.

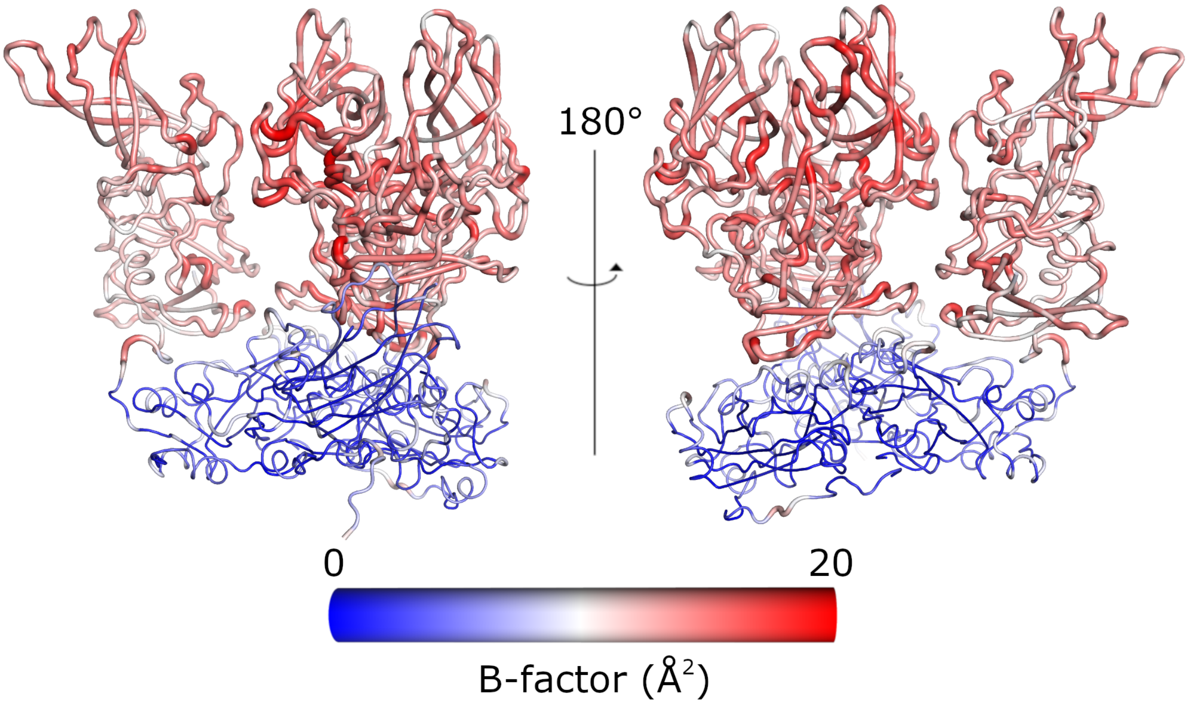


**Figure S4. B-factor 'putty' representation of wtMNV VP1 atomic model.** The atomic coordinates for wtMNV VP1 in each quasi-equivalent state, coloured by B-factor according to the scale shown. The backbone appears wider in parts of the model with higher B-factors.

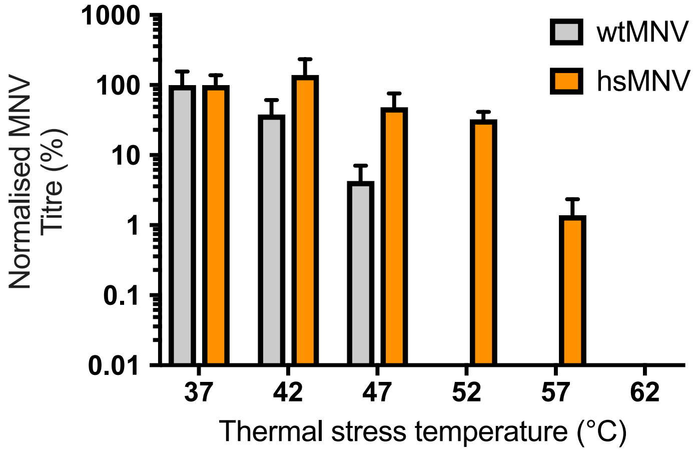


**Figure S5. Phenotypic validation of heat stable (hs)MNV.** wtMNV (grey) and hsMNV (orange) were incubated for 30 min at a range of different temperatures, then titred by $TCID_{50}$ assay on RAW264.7 cells ($n$ = 3 ± SEM).

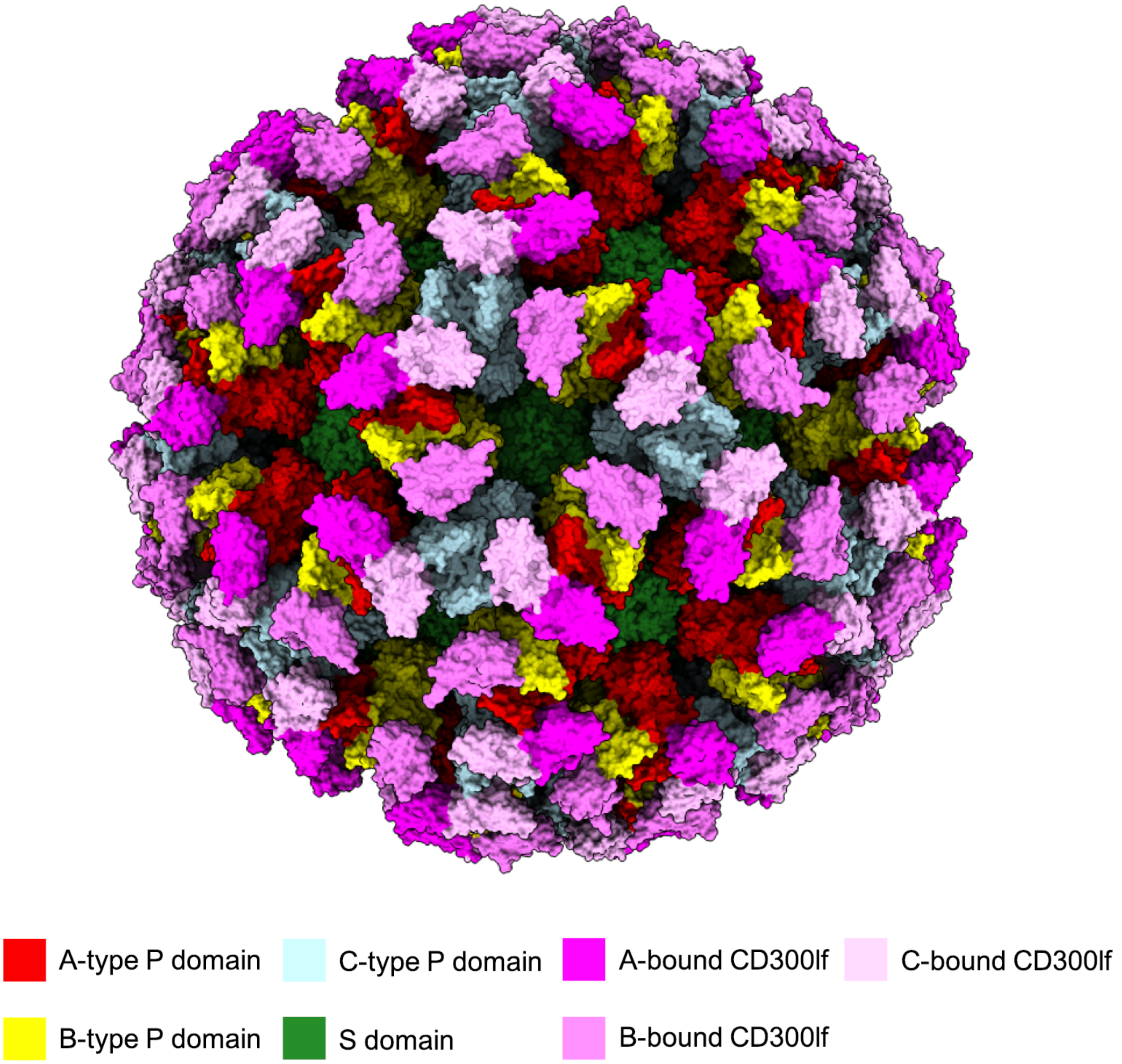


**Figure S6. MNV-CD300lf docking model.** The crystal structure of an MNV P domain dimer complexed with CD300lf (PDB: 6C6Q)[1] was rigid-body fitted into the wtMNV reconstruction along with the S domains of our VP1 atomic model to generate an asymmetric unit, which was symmetrised. Components of the model are coloured according to the scheme indicated and shown as molecular surfaces, viewed down the icosahedral three-fold axis.

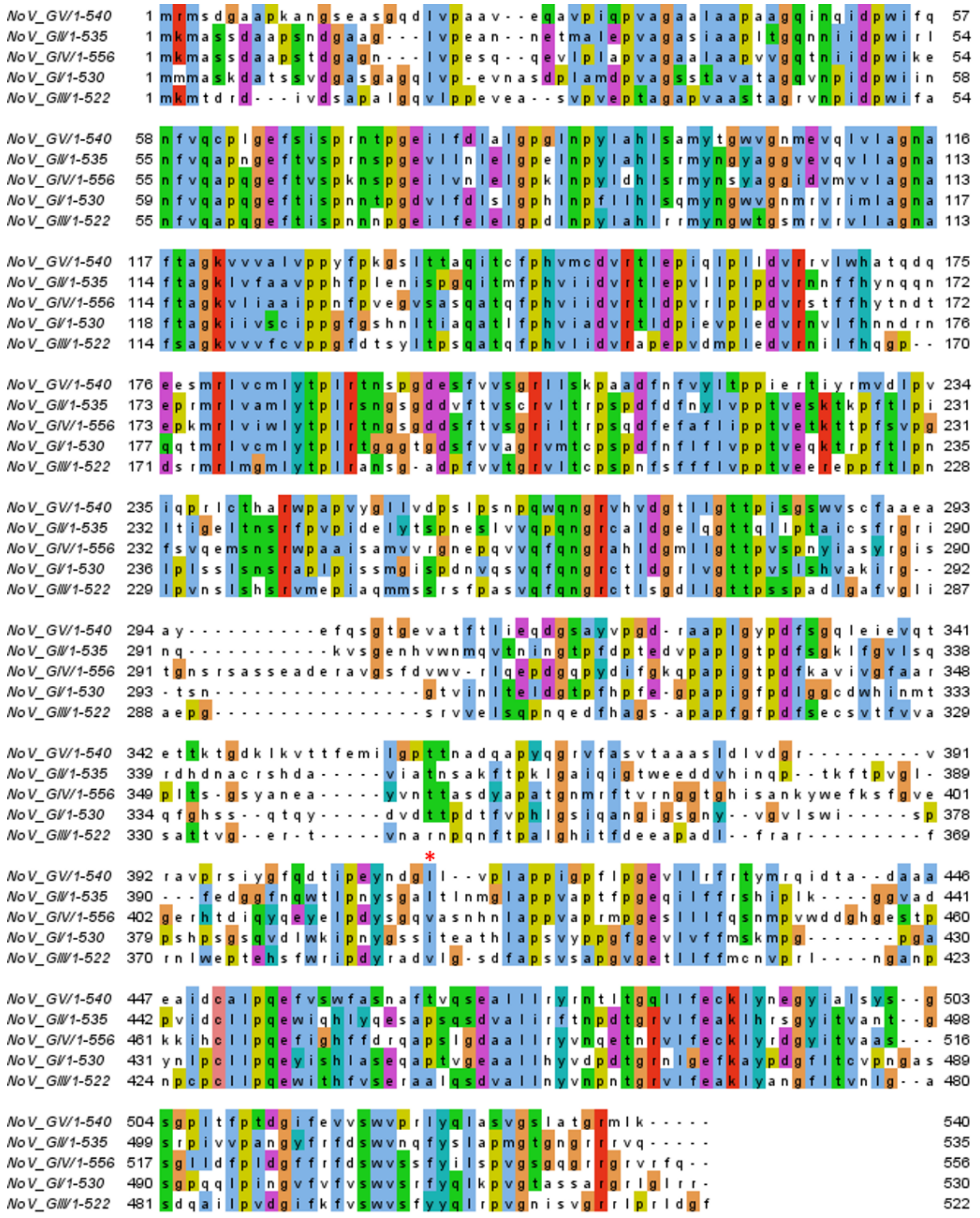

**Figure S7. Norovirus VP1 sequence alignment.** Alignment of VP1 sequences from norovirus genogroups GV (NCBI reference sequence: YP_720002.1), GI (NP_056821.2), GII (YP_009237898.1), GIII (YP_009237901.1) and GIV (YP_009237904.1). The MNV (GV) VP1 sequence is given in the top position. Sequences were aligned using Clustal Omega with default parameters[2–4], and are shown with the Clustal colouring scheme. MNV (GV) VP1 L412 is indicated by the red asterisk.

**Table S1:** Data collection and image processing parameters

| | wtMNV | hiMNV | hsMNV |
|---|---|---|---|
| Microscope | FEI Titan Krios | FEI Titan Krios | FEI Titan Krios |
| Camera | Falcon III | Falcon III | Falcon III |
| Voltage (kV) | 300 | 300 | 300 |
| Pixel size (Å) | 1.065 | 1.065 | 1.065 |
| Total dose ($e^-$/Å$^2$) | 59 | 59 | 64 |
| Number of frames | 59 | 59 | 59 |
| Defocus range (μm) | -0.5 to -2.9 | -0.5 to -2.9 | -0.7 to -3.0 |
| Number of micrographs | 13692 | 2620 | 7617 |
| Acquisition software | FEI EPU | FEI EPU | FEI EPU |
| Motion correction | MotionCor2 | MotionCor2 | MotionCor2 |
| CTF estimation | GCTF | GCTF | GCTF |
| Image processing | Relion 3.0 | Relion 3.0 | Relion 3.0 |
| Particles contributed | 7811 | 14266 | 35263 |
| B-factor | -129 | -136 | -153 |
| Resolution (FSC=0.143) | 3.1 | 2.9 | 3.1 |

**Table S2:** Model building and validation

| | |
|---|---|
| Model | **wtMNV VP1** |
| PDB ID | XXXX |
| Residues modelled<br>(from S domain homology model) | **A**: 20-191; 199-222<br>**B**: 17-221<br>**C**: 30-227 |
| Residues modelled<br>(from PDB: 6C6Q) | **A**: 228-531<br>**B**: 229-531<br>**C**: 228-531 |
| RMSD<br>*Bond lengths (Å)*<br>*Bond angles (°)* | <br>0.0139<br>1.42 |
| Validation<br>*All-atom clashscore*<br>*MolProbity score*<br>*Rotamer outliers (%)* | <br>5.43<br>1.66<br>0.16 |
| Ramachandran plot<br>*Favoured (%)*<br>*Allowed (%)*<br>*Outliers (%)* | <br>94.73<br>5.27<br>0.00 |

**Movie S1. Differences in P domain conformation between wtMNV from Katpally *et al.* (2010) and Snowden *et al.* (2019).** The crystal structure of an MNV VP1 P domain (PDB: 3LQ6) was rigid-body fitted into each quasi-equivalent position of the wtMNV EM density map from Katpally *et al.* (2010) (EMDB-7564) or the wtMNV map reported here, after low-pass filtering both to 8.0 Å resolution. The UCSF Chimera tool 'Morph Conformations' was used to switch between these two conformations. P domains are shown in orange. S domains (green) are also shown for clarity.

**Movie S2. hsMNV has 'twisted' AB-type P domains.** Morphing from the wtMNV (grey) to the hsMNV (orange) density maps highlights the change in AB-type P domain orientation. Maps are low-pass filtered to 8.0 Å and shown at 1.5 $\sigma$ with the back focal plane clipped to remove the S domains.

**Movie S3. An interface is disrupted between P domain dimers in hsMNV.** The atomic coordinates for wtMNV VP1 were separated into components (S domains, C-type P domain, and AB-type P domain dimer) and rigid-body fit then refined into the hsMNV density map. The change between these two conformations is shown, with P domains shown in orange, S domains in green, and the mutated residue (L412Q) highlighted in cyan. Density maps (wtMNV – grey, hsMNV – orange) are shown at 2 $\sigma$ after low-pass filtering to 5.0 Å resolution.

**Supplementary References**


1. Nelson, C. A. *et al.* Structural basis for murine norovirus engagement of bile acids and the CD300lf receptor. *Proc. Natl. Acad. Sci. USA* **115**, E9201–10 (2018).
2. Li, W. *et al.* The EMBL-EBI bioinformatics web and programmatic tools framework. *Nucleic Acids Res.* **43**, W580–4 (2015).
3. McWilliam, H. *et al.* Analysis Tool Web Services from the EMBL-EBI. *Nucleic Acids Res.* **41**, W597–600 (2013).
4. Sievers, F. *et al.* Fast, scalable generation of high-quality protein multiple sequence alignments using Clustal Omega. *Mol. Syst. Biol.* **7**, 539 (2011).
